# Reduced harmonic complexity of brain parenchymal cardiovascular pulse waveforms in Alzheimer’s disease

**DOI:** 10.1101/2025.02.26.640409

**Authors:** Anssi Koivula, Vilma Perkiömäki, Mika Aho, Aleksi Rasila, Valter Poltojainen, Vesa Korhonen, Matti Järvelä, Niko Huotari, Heta Helakari, Johanna Tuunanen, Lauri Raitamaa, Tommi Väyrynen, Johanna Kruger, Vesa Kiviniemi, Janne Kananen

## Abstract

Alzheimer’s disease (AD) is characterized by specific neuropathologies, and is associated with arterial wall β-amyloid accumulations, which lead to radiologically detectable amplitude increases and variable propagation speed of cardiovascular impulses in brain. In this study, we developed a fast frequency domain imaging method know as relative harmonic power of magnetic resonance encephalography (MREG_RHP_), aiming to investigate the configuration of the cardiovascular impulses independently of the mean magnetic resonance signal intensity and physiological impulse amplitude. In the initial analyses in healthy controls, we found that a wide 0.8 - 5Hz bandpass produced the most physiologically realistic cardiovascular waveforms. Whereas the data recorded in cerebrospinal fluid (CSF) from flip angle (FA) of 25° yielded up to 7-fold higher cardiac signal intensity as compared to FA of 5°, within the brain tissue recordings with FA of 5° were markedly more sensitive to cardiac waveform. We detected arterial impulses originating from major arteries and extending into the surrounding brain parenchyma, with simultaneous dampening of amplitude as a function of distance from source. Finally, we compared MREG_RHP_ results in 34 AD patients (mean age: 60.7±4.7 years; 53% female) against 29 controls (mean age: 56.9±7.9 years; 66% female). We show that the harmonic power of cardiovascular brain pulses is significantly reduced in cortical frontoparietal areas of AD patients, indicating monotonous impulse patterns colocalizing with the previously reported areas of increased impulse propagation speed. In conclusion, the MREG_RHP_ offers a fast Fourier transform (FFT)-based method to non-invasively quantify and locate human arterial blood vessel wall pathology.

## 1. Introduction

Since the first clinical and histopathological description of Alzheimer’s disease (AD), its characteristic neurodegeneration has been linked to aggregation of abnormally-folded proteins in the brain parenchyma. Recent research has implicated, failure of a brain-wide solute clearance mechanism mediated by cardiovascular pulsation of perivascular cerebrospinal fluid (CSF) known as the glymphatic pathway, to the pathogenesis of AD (Nedergaard and Goldman, 2020). Whereas most preclinical studies of glymphatic function have used invasive fluorescent tracer microscopy in rodent models, corresponding studies in healthy humans and patients with normal pressure hydrocephalus have used magnetic resonance imaging (MRI) after intrathecal gadolinium injection to track the kinetics of tracer distribution within the CSF (Eide and Ringstad, 2015; Ringstad et al., 2018; Ringstad and Eide, 2020). Due to its invasive nature, it is challenging to apply this methodologically elegant approach in AD populations, which calls for the development of novel and non-invasive techniques.

Functional magnetic resonance imaging (fMRI) has been successfully applied to investigate changes in the slow synchrony of brain network connections with respect to declining cognitive function in AD patients (Greicius et al., 2004). Such methods have revealed increased brain signal variability in AD (Makedonov et al., 2016). Furthermore, increased coefficient of variation (CV, also known as the relative signal standard deviation) in blood oxygenation level dependent (BOLD) signals has been linked to cerebrovascular pathology in non-demented individuals (Jahanian et al., 2014) and has emerged as a promising biomarker of AD-related neuronal degeneration (Scarapicchia et al., 2018; Tuovinen et al., 2020). More recently, faster fMRI sequences with up to 5 Hz frequency resolution have been employed to obtain critically sampled T2* weighted brain signals, which reflect classical BOLD signal from veins, as well as faster-propagating arterial spin phase/steady state free spin precession (SSFP) effects arising from cardiovascular impulses (Von Schulthess and Higgins, 1985). Fast fMRI sampling, known as magnetic resonance encephalography (MREG), has enabled the detection of altered cardiovascular brain pulsations in AD without aliasing of different signal components (Rajna et al., 2021, 2019; Tuovinen et al., 2020); the data indicate increased amplitude variance and propagation speed of cardiovascular pulses in frontoparietal areas, and reversed propagation in hippocampal regions, which are involved in the encoding of memory.

Factors relating to differing instrumentation, vendors, and environment hinder the standardized evaluation of fMRI signal amplitude due to fluctuations in mean signal levels and effects on overall signal gain. The consequent effects on statistical metrics such as standard deviation (SD) and CV can potentially lead to misinterpretation of fMRI. This limitation calls for development of methods that are agnostic to signal amplitude *per se*. Furthermore, the higher spectral harmonics of cardiovascular pulsations can convey important information regarding CSF flow dynamics (Yildiz et al., 2022, 2017). A method that effectively utilizes harmonic spectral features and is agnostic to mean signal level is known as total harmonic distortion, which is a metric commonly used in electrical engineering to quantify the power of spectral harmonic signal components with respect to the principal frequency power. In general, any system with low harmonic distortion suffers from unintentional interference from other inputs, while a system with high harmonic distortion is less vulnerable to interference.

In this article we introduce an amplitude agnostic method for estimating waveform complexity in brain, based on quantifying the relative harmonic power of included frequency components (MREG_RHP_) in the wideband cardiovascular fMRI signal. In our approach, we first optimized the frequency band width for quantifying the MREG_RHP_, and next evaluated two different flip angles (FAs; 25° and 5°) for optimizing the detection of cardiovascular impulses. Finally, we tested the hypothesis that pathological changes in the physiological signal in AD brain could be detectable in the relative cardiac impulse waveform shape. The analysis of MREG_RHP_ differences between patients with AD and healthy controls confirmed our hypothesis by revealing significant differences in harmonic features of the cardiovascular impulse shape in AD.

## 2. Materials and methods

### 2.1 Study population

The study was performed in accordance with the Helsinki declaration and current good clinical practice guidelines, with approval by the Northern Ostrobothnia Ethical Committee (53 & 118/2012). Patients with AD (n = 34) and age-matched controls (CON; n = 29) were recruited during 2016 – 2019, after their provision of written informed consent. (Table 1).

**Table 1.**

| **Characteristic** | **AD (n=34)** | **Controls (n=29)** | **p-value** |
| --- | --- | --- | --- |
| Females | 18 (52.9%) | 19 (65.5%) | 0.44 |
| Males | 16 (47.1%) | 10 (34.5%) |  |
| Mean age when imaged (years, SD) | 60.7 (4.7) | 56.9 (7.9) | 0.022 |
| Mini-Mental State Examination (SD) | 22.1 (6.0) | not applicable |  |
| Disease duration (years, SD) | 3.8 (3.4)* | not applicable |  |
| Mean heartrate (SD) | 64.1 BPM (11.9) | 65.8 bpm (9.1) | 0.517 |
| Mean arterial blood pressure (SD) | 151.7/81.6 mmHg<br>(22.2/9.7)** | 145.6/82.9 mmHg<br>(22.7/10.5)*** | 0.340/0.654 |

\* data missing from 1 patient
\*\*data missing from 8 patients
\*\*\*data missing from 5 controls

### 2.2 Imaging and pre-processing

All subjects were scanned with a Siemens 3T SKYRA scanner with a 32-channel head coil. Subjects were scanned with a MREG sequence, which under samples the k-space using a 3D single shot-stack of spirals (SOS) trajectory to reach a sampling rate of 10Hz (i.e., 10 whole brain datasets per second), thereby allowing critical imaging of physiological pulsations (Assländer et al., 2013). The SOS gathers the k-space with spiral in/out repeating in every other turn continuously in the positive z-direction to minimize off-resonance artifacts from the air sinuses (for more details, c.f. Assländer et al., 2013). The sequence enabled scanning of the whole brain with a voxel size of 3×3×3 mm^3^ (TR=100 ms, TE=36 ms, flip angle (FA)= 25° or 5°, 3D matrix=64^3^, FOV=192 mm).

The magnetization spoiling gradient between scans was set to 0.1 to avoid stimulated echo drifts masking the signal, while retaining sensitivity to the physiological pulsations. Individual subject MREG data were reconstructed by L2-Tikhonov regularization with λ = 0.1, and the L-curve method was used to determine the regularization parameter (Hugger et al., 2011). The resulting effective spatial resolution was 4 mm. Image reconstruction included a dynamic off-resonance in the k-space (DORK) method, which corrected for scanner warming and respiration-induced dynamic B_o_-field changes before preprocessing (Zahneisen et al., 2011). Supportive cardiorespiratory data were collected using an MRI-compatible multimodal neuroimaging system (Korhonen et al., 2014). Physiological measurements were recorded with an anesthesia monitor, with heart and respiratory rates identically matching the MREG-data in time and frequency domains, as previously shown (Järvelä et al., 2022; Kananen et al., 2022; Tuovinen et al., 2020).

FSL BET was used for T1 anatomical brain extraction (fractional intensity = 0.25, threshold gradient = 0.22, neck and bias-field correction) for further processing of the MREG data. Next, the data were examined for outliers with AFNI *3dToutcount* to reveal potential spiking of data caused by coarse subject movement during scanning. A single continuous time-sequence per subject (with the least possible amount of coarse movement) was then extracted from the original scan data for further pre-processing. These shorter sections were typically 2701 time points (270 seconds at 10 Hz) in length. Next the dataset was despiked with AFNI *3dDespike* using *–NEW25* and *-localedit* options and preprocessed using a typical FSL pipeline (Beckmann and Smith, 2004), including high-pass filtering (cut-off frequency of 0.008 Hz, i.e. 125 s), motion correction (FSL MCFLIRT, Jenkinson et al., 2002), and transformation into MNI152-space.

### 2.3 Filtering of the main cardiovascular waveform and harmonic components

The 10 Hz MREG spectrum range is 0 - 5 Hz according to the Nyquist-Shannon sampling theorem, and thus allows the detection of up to four harmonics of a < 60 bpm (1 Hz) cardiac cycle (Figure 1). AFNI *3dTproject* was used for filtering the data in three ways: narrow band around the cardiac principal frequency, extended (encompassing the principal cardiac peak +1 harmonic) and wide bandwidths (from the principal cardiac peak to 5Hz) having maximal harmonic coverage, as shown in Figure 1A. Cut-off frequencies were selected manually for each subject based on their recorded respiratory and cardiac rate upper and lower limits, thus ensuring sufficient coverage of their physiological variations. Respiratory bands and lower heterodyne peaks were cut from the high pass filtered signals.

**Figure 1.**
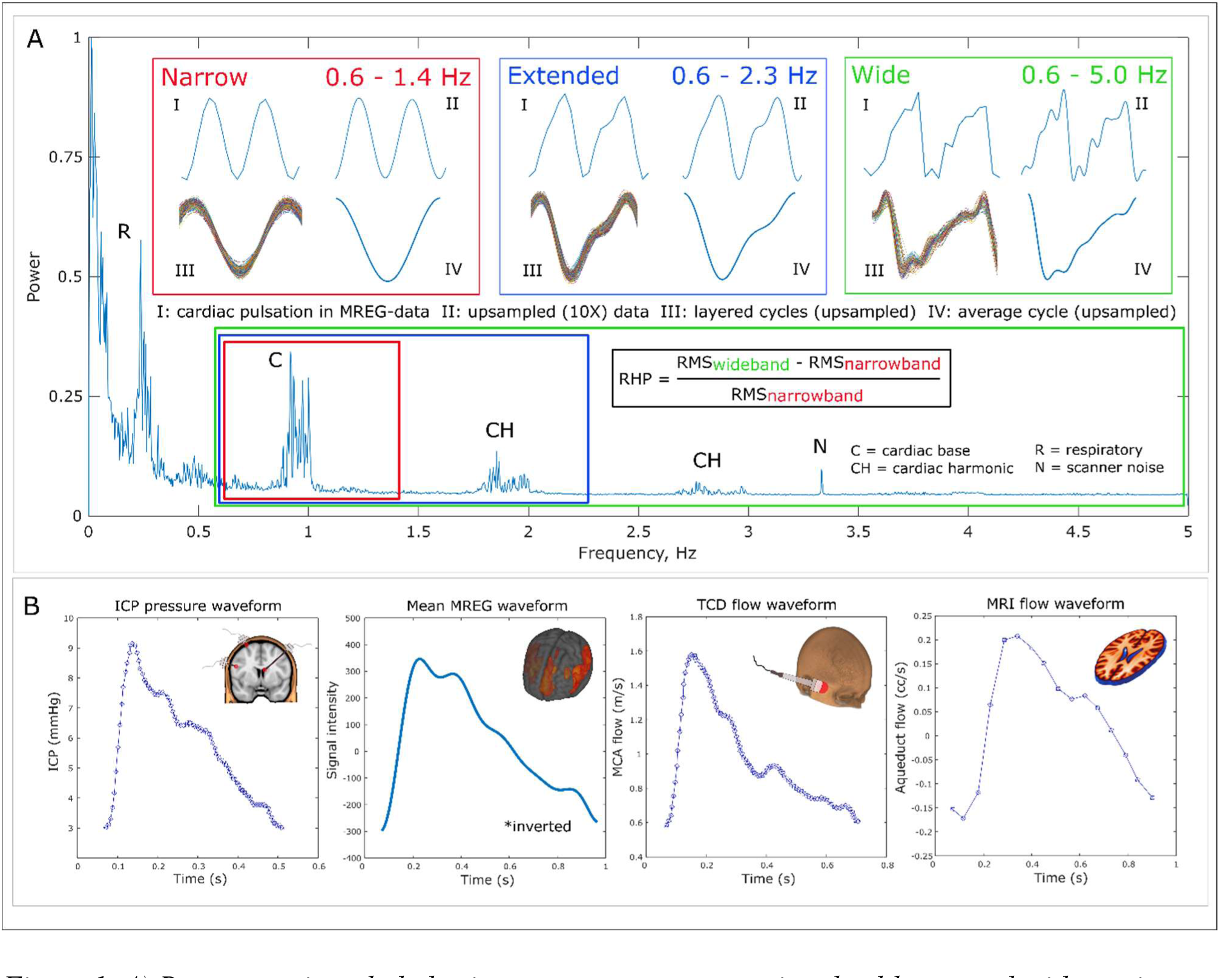
A) Representative whole brain mean power spectrum in a healthy control with respiratory and cardiac peaks (including principal and harmonic frequencies) superimposed over the typical 1/f trend curve. The narrow (red) extended (blue) and wide (green) filtering bandwidths reveal how the signal waveform transforms from a pure sinusoid to had more physiologically realistic features upon inclusion of more harmonic components. Relative harmonic power (RHP) is a measure for waveform complexity based on the relation of the harmonic signal component power to the principal cardiac component power. B) Example waveforms of four different modalities measuring intra-cranial pulsations. Intracranial pressure (ICP), transcranial doppler (TCD) and magnetic resonance imaging (MRI) waveforms are adapted from Wagshul et al. (2011). Observe the homologous features with local MREG, including distinct systolic/diastolic phases and the dicrotic notch.

### 2.4 Waveform complexity estimation with MREG_RHP_

We estimated the cardiovascular waveform complexity by calculating the ratio of the harmonic signal component power against the principal cardiac component power. Consequently, the measurement is designated *Relative Harmonic Power of MREG* (MREG_RHP_). The basic principle is analogous with total harmonic distortion, a measurement commonly used in electrical engineering used to quantify the power of harmonic signal components (Blagouchine and Moreau, 2011). A previous study utilizing multi-slab echo-volumar imaging with comparable whole brain coverage and a TR of 136 ms (Posse et al., 2013), used a somewhat similar, but inverse ratio, with only the second harmonic component, to evaluate cardiac-related pulsations in different areas of the brain.

### 2.5 Selection of test voxels

Test and reference voxels were needed for analyzing MREG_RHP_ waveforms. The selection of these voxels was based on well-established knowledge of the anatomical locations of the major cerebral blood vessels: the anterior cerebral artery (ACA), the middle cerebral artery (MCA) and the superior sagittal sinus vein (SSAG). In addition, brain-wide pulsatility maps were created by calculating root mean square (RMS) values separately for each timeseries voxel-by-voxel. These maps revealed brain regions and single voxels with the highest pulsatility, thus guiding the selection process of voxels depicting ACA, MCA and SSAG.

### 2.6 Statistical methods and tests

The mean and standard deviation values of demographic data were calculated in SPSS version. 26. The p-value for differences in sex distributions represents an exact two-sided Pearson Chi-Square test. For systolic and diastolic blood pressure, heart rate, age, and Mini-Mental State Examination (MMSE), the p-values represent results from two-sample t-tests. Population mean MREG_RHP_ brain maps for the AD and CON groups were compared with FSL *randomise* with threshold-free cluster enhancement and family-wise error rate (FWER, p < 0.05) correction and 5000 permutations.

## 3. Results

### 3.1 Study group demographics

The mean (SD) age of the AD group (60.7±4.7 years) was significantly 7% higher than in the CON group (56.9±7.9 years) (p = 0.022, Table 1). Other parameters including proportion of females, heart rate, systolic and diastolic blood pressure, and MMSE score were similar between groups (p > 0.05, Table 1).

### 3.2 Cardiovascular pulse waveforms in the brain

After preprocessing of the fast fMRI data, we calculated whole brain power spectrum with fast Fourier transform (FFT, Figure 1A). Figure 1 illustrates how the narrow filtering option (red) contains only the narrow principal cardiac frequency, such that the waveform has a sinusoidal appearance. The extended filtering option (blue) contains cumulatively the 1st harmonic frequency component, such that the time domain signal has the classical dicrotic notch shape, as illustrated in the arterial signal. The wide filtering option (green) reaches up to 5 Hz, including the 1^st^ - 4^th^ harmonics in subjects with < 1 Hz (>60 bpm) heart rates. This procedure transforms the waveform into a more physiologically realistic shape with a sharp systolic signal drop followed by a slower diastolic phase return with a classic dicrotic notch (Figure 1B); the resultant waveform has more subtle intrinsic signal complexity, being comparable to invasive analogue intracranial pressure measures and fast sampling transcranial doppler ultrasound, but superior to conventional gated phase MRI data.

### 3.3 Morphology of cardiovascular pulsations in brain tissue

We continued our analysis using the more physiological wideband cardiac-filtered high-pass signal, thereby detecting high-amplitude pulsations (based on RMS maps) in voxels depicting major cerebral blood vessels and spaces filled with CSF. Examples of continuous MREG-waveforms from different parts of the brain are presented in Figure 2A. The two high-amplitude waveforms from MCA (red) and SSAG (blue) appear to be well synchronized in time, but intriguingly are in opposite phase. In grey matter (cyan), the waveform shape is more symmetrical, and the signal amplitude attenuates in relation to the penetration depth of the cardiac impulse into brain tissue. In white matter (green) the signal amplitude is even further attenuated and the periodic pattern barely visible using the same amplitude scaling. More advanced analysis in time-domain revealed interconnectedness and an evident spatially heterogeneous distribution of these waveform types (Figure 2).

**Figure 2.**
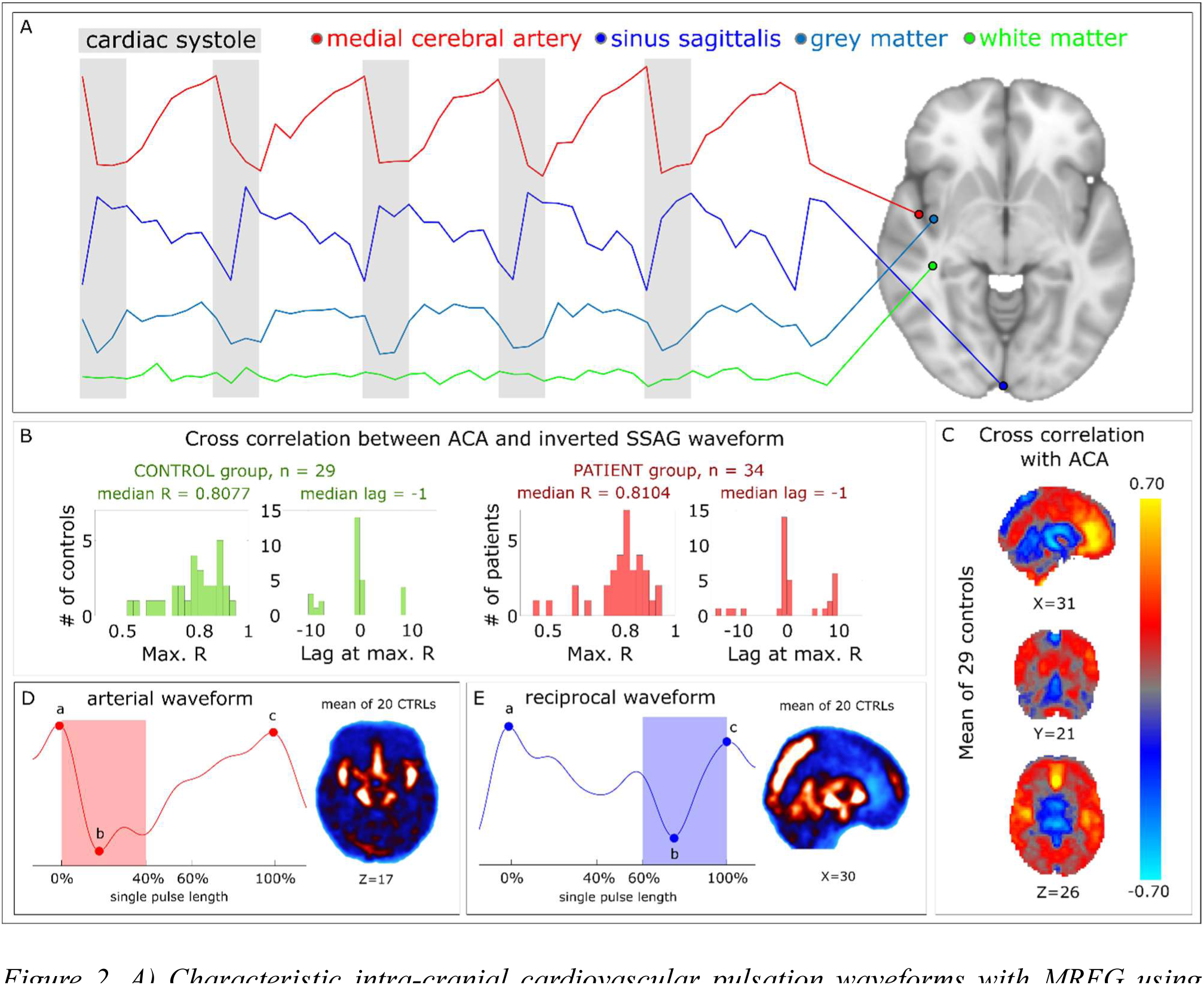
A) Characteristic intra-cranial cardiovascular pulsation waveforms with MREG using wideband filtering in a representative healthy control. During systole (highlighted gray background), signal intensity rapidly drops in large cerebral arteries (red) and increases in large veins (blue), creating a high-amplitude sharp-edged pattern. In the insular grey matter (cyan), amplitude decreases, and the waveform becomes more symmetrical with a preserved periodic nature. In white matter (green) signal amplitude is markedly lower and the periodic pattern barely visible using the same intensity scale. B) There was a strong correlation between arterial signal from ACA and inverted SSAG signal, and the arterial signal preceded the sagittal sinus waveform by a median of 0.1 seconds (based on cross correlation phase lag) in both study groups. C) Brain-wide cross correlation with an arterial signal from ACA highlights arterial and cortical structures with positive correlation, in contrast with the negatively correlating CSF and venous structures. D) In a tentative complimentary waveforms analysis for 20 controls, the arterial waveforms the nadirs were located within initial 40% of the cardiac pulse length (red shadowing). Spatially these cardiovascular waveforms were strongly present in major arteries of the circle of Willis. E) The reciprocal venous waveforms related to cardiovascular pulses had their nadirs in the latter 60 - 100% of the pulse length (venous shadowing). These reciprocal waveforms were found in large veins, but also in central CSF spaces.

We next performed voxel-by-voxel temporal analysis to reveal spatial distributions of the two high-amplitude, opposite-phased waveform types identified in MCA and SSAG. Control group mean results and basic categorization principles for pulse-to-pulse analysis are illustrated in Figure 2B-E. Henceforth, we refer to the MREG signal waveform type characterized by a sudden drop in signal intensity as being *arterial*, as it is prominent in the major cerebral arterial branches of the circle of Willis (Kiviniemi et al., 2016; Posse et al., 2013, Rajna et al., 2021). Respectively, we designate the second waveform type as *reciprocal*, since it is opposite-phased to the *arterial* waveform type. The reciprocal waveform is prominent in certain parts of the intra-cranial venous sinuses, and is also present in basal CSF spaces and ventricles, such that calling it *venous* would be only partially correct. When, for example, a venous signal from the sagittal sinus is inverted and plotted over a signal from the anterior cerebral artery, the similarity and synchronous behavior of the two waveforms is evident (Figure 2D-E). In both subject groups, the median coefficients for cross-correlation between the arterial and inverted venous waveform over the whole timeseries were approximately 0.8, indicating a strong correlation (Figure 2B). Maximal correlation scores were achieved with a median lag of minus one timepoint (0.1s), when the permitted lag range was set to ± 1.5 s (i.e. 15 timepoints, Figure 2B). In other words, the inverted venous signal lags behind the arterial waveform by 0.1 s.

The simple categorization (arterial/venous) principle is shown in Figure 2D-E. In brief, each nadir (point b) is coupled with a preceding peak (point a) and a following peak (point c), which together define the wavelength of a particular pulse. In *arterial* waveforms the nadirs are located between the initial 0 - 40% of the cardiac pulse (Figure 2D, red shadowing), after which the signal returns to baseline. At the same time, in the *reciprocal* (venous) waveforms the signal initially increases and then drops to a nadir located in the final 40 % of the cardiac pulse (Figure 2E, blue shadowing). The strength score of a waveform type is defined by the number of detected arterial or venous pulses in each timeseries of a given duration. By application of these principles, we obtained reliable positioning of major arterial and venous structures.

Brain-wide cross correlation with the strongest and most steady arterial reference signal (Rajna et al., 2021), which is from the ACA (Figure 2C), can be used to distinguish arterial from venous behavior in parenchymal voxels containing weaker amplitude or otherwise spurious or ambiguous signals (Figure 2C-E). In these figures, positive correlation values indicate similarity with the arterial waveform and the highest correlations match anatomical locations of the main cerebral arteries and cortical grey matter supplied by the ACA. Conversely, areas with negative correlation values match mainly with the sagittal and cavernous sinus and ventricles. The pulsations in these CSF-filled areas are opposite-phased to the arterial waveforms. Interestingly, parts of the cerebellum and the adjacent subratentorial CSF space in proximity of sinus rectus also manifest opposite-phased relative to the ACA.

### 3.4 MREG flip angle and signal waveforms

We have previously used two different flip angles (FA 5° and FA 25°) for analyzing AD brain pulsations (Rajna et al., 2021; Tuovinen et al., 2020). Insofar as flip angle is a key imaging parameter for physiological “noise” in fMRI signal (Gonzalez-Castillo et al., 2011), we investigated the differing ability of these two flip angles for detecting arterial signal waveform characteristics and their spatial distributions. The results for control subject data revealed major differences between FA 5° and FA 25° in their wideband cardiac signal variabilities (Figure 3). Indeed, the group mean cardiovascular signal intensity with data in all CSF spaces was up to seven-fold higher with FA 25° data compared to FA 5°, reflecting its greater sensitivity to physiological (i.e. respiratory) signal sources apt to increase spin history effects. On the other hand, the detection of arterial waveforms in brain tissue (using the criteria presented in Figure 2D) was markedly more accurate with FA 5° data compared to FA 25° data. This conclusion is substantiated by comparing the detection results with a time-of-flight reference blood vessel atlas (Viviani, 2016) and the brain-wide cardiovascular correlation structure presented in Figure 2C. This result led to us to place our focus on FA 5° data in the wide spectral range analyses.

**Figure 3.**
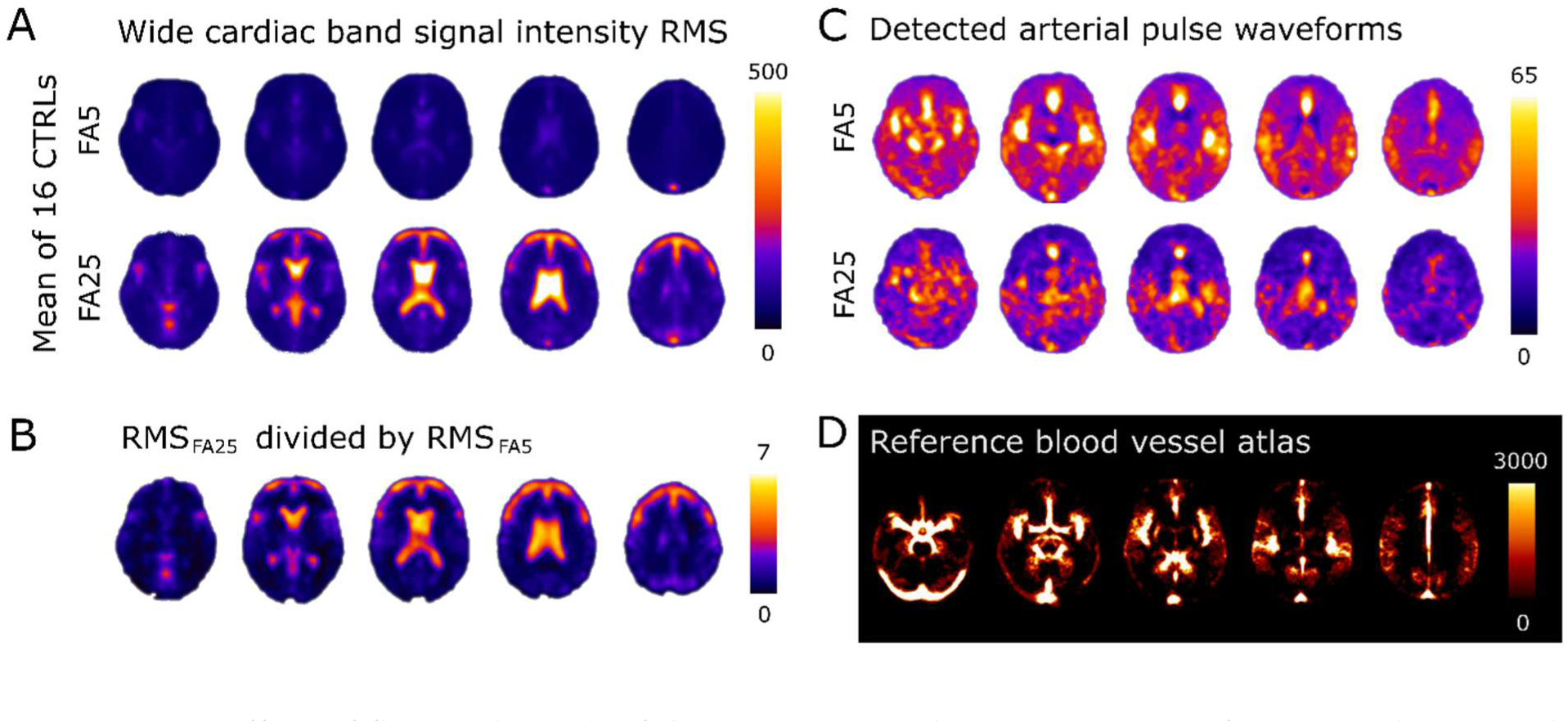
A) Effect of flip angle (FA) of the MREG signal in comparison of FA 5° and FA 25° data in the subset of (n=16) controls having both FA 5° and FA 25° data. B) Group mean signal intensity with FA 25° data in the ventricles and other CSF spaces was up to seven times that of FA 5° data. C) Regardless of the higher signal variability of FA 25° data, detection of arterial waveforms was spatially more accurate with FA 5° data when compared with FA 25°data. D) A reference blood vessel atlas (Viviani, 2016).

### 3.5 Reduced MREG_RHP_ waveform complexity in Alzheimer’s disease

MREG_RHP_ estimates the complexity in the cardiovascular waveform using the power of harmonic signal components in relation to that of the principal cardiac frequency. Group-level mean results for FA 5° data using MREG_RHP_ are shown in Figure 4A, along with axial slices for visual comparison. In the two subject groups, the MREG_RHP_ scores were similarly low, with little signal complexity in areas matching anatomically with major cerebral blood vessels and CSF spaces. In contrast, the highest MREG_RHP_ scores were found mainly in white matter for both groups. Statistically significant group differences (p < 0.01, FSL *randomise*, FWER-corrected) occurred over large areas of the brain parenchyma, including white matter, cortex, the basal ganglia, and at the edges of the lateral ventricles. However, the signals in large blood vessels and central parts of ventricles did not differ between groups.

**Figure 4.**
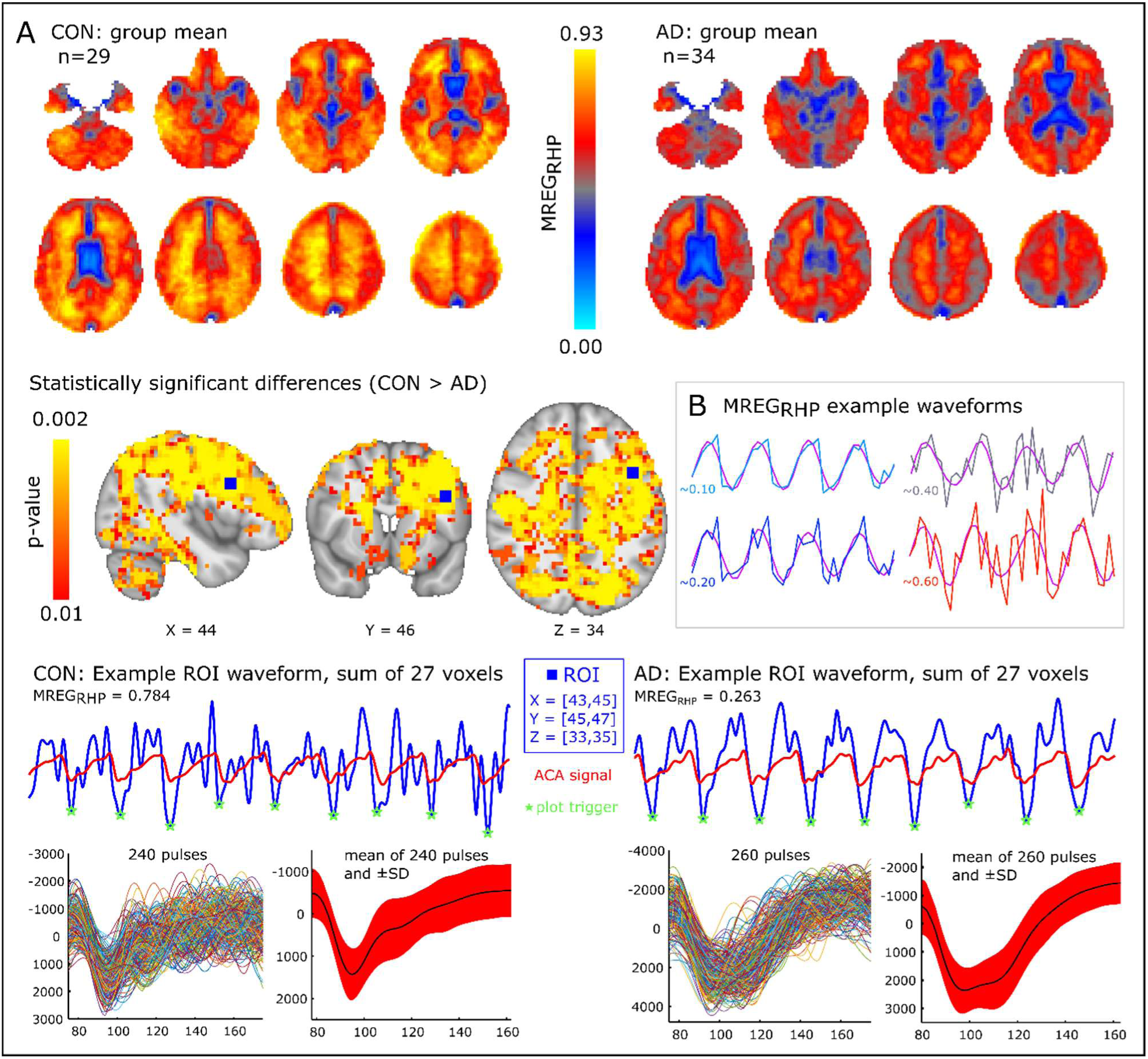
A) Based on mean MREG_RHP_ results, intra-cranial cardiovascular waveform complexity is decreased in Alzheimer’s disease (AD). Statistically significant differences between AD and healthy controls (CON) groups were found in the brain parenchyma, but not in major blood vessels or the ventricles. Region-of-interest (ROI) waveforms for representative single CON and AD subjects were formed by summing together 27 wide cardiac band timeseries from a 3×3×3 voxel area. The ROI center point (X=44, Y=46, Z=34) was selected based on the smallest p-value in group-wise comparison of MREG_RHP_. Nine cardiac cycles, (upsampled) of the ROI waveforms are shown. Waveforms from the anterior cerebral artery (ACA) are included as a reference for cardiac output. Individual pulses were then extracted from this ROI over the whole timeseries, and then aligned and projected on top of the others. Mean (SD) pulse waveforms evidently reflect the group differences in signal complexity (0.784 vs. 0.263) and MREG_RHP_ between the CON subject and the AD subject respectively. B) Four modelled example waveforms further substantiate the nature of signal complexity with simultaneously increasing levels of MREG_RHP_. We note the differences between principal cardiac frequency waveforms (purple color) and the more complex wideband waveforms.

Figure 4B illustrates representative single voxel waveforms (extracted from one individual in the CON group) with levels of MREG_RHP_ ranging from 0.10 to 0.90. These examples further substantiate that the principal cardiac frequency waveform (narrow filtering option, sinusoidal-like in purple color) relates with the wideband signal with all available harmonic components as a function of MREG_RHP_. In the instance of low MREG_RHP_ in the AD group, the cardiac waveform exhibited a nearly sinusoidal pattern, with a loss of mainly diastolic signal complexity, notwithstanding the wide spectral filtering (Figure 4A), while the waveforms were more complex in brain tissue of the AD group, with prominent spiking solely during diastole, thereby sensitively capturing the pulse wave reflections from vasoactive arteries and arterioles (Figure 4A).

## 4. Discussion

In this paper we optimized brain-wide detection of arterial pulse waveform patterns using noninvasive ultra-fast MREG technology to quantify and map AD-related changes in arterial pulsation. We first optimized the FFT filtering bandwidth to obtain realistic cardiovascular impulse waveforms with plausible physiological characteristics. We then determined that the lower flip angle (FA 5°) performed better than FA 25° in detecting intraparenchymal arterial pulse waveforms, due to the lesser effects of respiratory noise. After analyzing the wideband cardiovascular pulse waveforms of the (FA 5°) MREG data, we found that the cardiovascular impulses in the brain tissue were monotonic and of lesser diastolic waveform complexity in the AD group as compared to the controls, who displayed greater complexity in the diastolic part of the arterial waveform.

### 4.1 Improved cardiac waveform detection

Intracranial cardiovascular pulsation waveforms are by no means sinusoidal, as our present data and other studies demonstrate (Carrera et al., 2010; Claassen et al., 2021; Wagshul et al., 2011). Moreover, previous investigations in cerebrovascular lesions and between different brain regions have already identified the significance of morphological changes in these pulsations (Posse et al., 2013). Contrary to conventional slowly sampled fMRI sequences that are hindered by aliasing of physiological signals (Huotari et al., 2019), we utilized ultrafast MREG for precise imaging of cardiovascular impulse features up to 5 Hz. This approach enabled clear-cut detection of a sharp systolic signal drop followed by a slower diastolic phase over the cardiac cycle in the vicinity of arteries in accord with earlier results (Claassen et al., 2021; Wagshul et al., 2011). The MREG technique’s augmented measurement points in the time domain and detection of harmonic signal components in the frequency domain enabled extensive brain-wide analyses of waveform shape, timing, and phase.

By simply measuring how the induced signal peaks and nadirs in each MREG timeseries are organized temporally in relation to each other, we were able to identify opposite-phase *arterial* and *reciprocal* waveforms in brain voxels with high signal amplitude. This alone sufficed to spatially map the large arteries and brain cortex from fMRI data, while the opposed phase signal mapped venous sinuses and CSF spaces. Unambiguous sorting of voxels into either *arterial* or *reciprocal* categories is problematic in the brain parenchyma, especially at the capillary level, and in white matter, where signal amplitudes are lower and waveform shapes more ambiguous (Santisakultarm et al., 2012). Because of this limitation, we utilized brain-wide cross-correlation with a strong *arterial* reference, i.e. anterior cerebral artery, to reveal pulsation phasing also in more weakly pulsating parenchymal voxels. The resulting distribution of positively (arteries, cortex) and negatively (cortical sulci, veins, ventricles, and aqueducts) correlating areas proved to match anatomy to structural MRI. Interestingly, however, pulsation in wide areas of the cerebellum, around sinus rectus, and in the cerebral aqueduct had opposed phase to the arterial references, including the basilar artery.

### 4.2 Cardiovascular pulsation effects on T2* weighted signal

The bi-phased *arterial* and *reciprocal* cardiovascular signals derive from basic concepts of cardiovascular physiology and MRI physics. The fast impulse-like changes in signal intensity during systole are caused by a rapid rise in arterial blood pressure and consequently accelerated blood flow (Duyn, 1997; Kiviniemi et al., 2016; Posse et al., 2013; Von Schulthess and Higgins, 1985). During the cardiovascular accelerative systolic impulse, the (peri)*arterial water spins* diphase and furthermore transiently lose their steady SSFP coherence due to the blood flow impulse, such that the MR signal intensity rapidly decreases, due to a wave propagating through neighboring brain water molecules (Duyn, 1997; Von Schulthess and Higgins, 1985). During diastole, the water ^1^H spins return to a baseline steady state, as the pressure wave passes and dissipates until the next impulse arrival.

Importantly, the arterial impulse effect is not a susceptibility effect like the BOLD contrast in veins but rather represents a temporary spin phase/SSFP effect, which precedes the BOLD effect in the local arteries (Huotari et al., 2022). In veins, the signal change is of opposite direction to the arterial signal, instead showing a rapid increase in signal intensity in major veins like the sagittal sinus. The increased venous signal follows the drop in arterial impulse with a median time delay of 0.1 s, which is in good agreement with the 0.106-0.171 s mean lag detected on healthy elderly controls by Rivera-Rivera (2017) with flow MRI. This phase shift most likely arises from a compensatory outflow of venous blood to remove an equal volume of blood (at 60 bpm with 1000 ml per minute of cerebral blood flow = 17 ml/beat) with each arterial impulse, following the Monro-Kellie doctrine (Claassen et al., 2021; Wagshul et al., 2011). The resulting reduction in venous deoxygenated blood volume leads to an increase in BOLD signal intensity, in analogy to the venous volume effects after hyperemia induced by neuronal activation. Moreover, the induced increase in the perivenous CSF water content, which replaces the loss of venous cerebral blood volume, further contributes to the increase in MRI signal as a T2 effect. Furthermore, the venous signal seems to depend upon its orientation relative to the spiral gradient trajectories of the 3D MREG sequence with respect to the scanneŕs B0-field. In our data, the signal amplitude was lower in directions perpendicular to the external field, such as in the transverse sinus, and higher in the posterior parts of the sagittal sinus, which has a cranio-caudal orientation. This is in keeping with an earlier report that BOLD signal intensity changes are strongest in microvascular structures running in parallel with B0 fields, and of lowest variability in perpendicular directions (Báez-Yánez et al., 2017).

### 4.3 Flip angle - sensitivity to physiological signal

According to a previous BOLD-signal sensitivity study (Gonzalez-Castillo et al., 2011), low flip angles impart lesser physiological “noise” and consequently improved tissue contrast. Present complementary findings indicate detectability of cardiovascular signal features is also highly dependent on the flip-angle. Signal intensity in CSF spaces was up to seven-fold higher with FA 25° compared to FA 5° data. However, signal variability in parenchymal voxels was far less affected by flip-angle, likely due to their more heterogeneous mix of tissue types interspersed with cortical CSF spaces, and inclusion of signals arising from arteries and veins distinct different flow and oxygenation characteristics. In addition, FA 5° gave a more accurate detection of arterial waveforms. These observations together imply that a lower flip-angle can suppress spurious physiological signal sources mediated by the flow of CSF, a deduction that is line with the studies mentioned above. Hence, interpretations of brain physiology and pathology studies must account for the impact of flip angle.

### 4.4 Complexity of pulsation waveforms in AD

The various MR signal metrics are affected by vendor, scanner type, field strength and homogeneity, signal gain, and relative mean intensity levels. To minimize the effects of these potential confounding factors, we developed the MREG_RHP_ method for evaluating brain-wide cardiovascular pulsation biometrics based on the waveform shape, rather than intensity. We then obtain an indirect estimation of waveform complexity based on the ratio between the power of harmonic signal components to the principal cardiac frequency power. Our method is thus indifferent to pulsation phasing, and can be applied to parenchymal voxels with low signal amplitude, in which the more conventional estimates such as pulsatility index, resistance index, and AUC cannot be reliably determined.

Given its evaluation of pulse shape irrespective of amplitude the RHP approach may inadvertently lose amplitude-based information relevant to brain physiology. On the other hand, MREG_RHP_ is less vulnerable to potential sources of error originating from imaging parameters or any other technical feature affecting signal gain. For example, were a timeseries amplitude increased/multiplied by a factor of two due to any non-physiological reason, its SD would also increase twofold, whereas the waveform shape as such and MREG_RHP_ results would be unaffected by the signal doubling.

MREG_RHP_ revealed significant differences between AD patients and controls extending across the entire brain, but excluding large blood vessels and centroids of the lateral ventricles. In general, complexity of the pulse waveform in the brain parenchyma was significantly lower, i.e more monotonic in the AD group. Another study with conventional rs-fMRI data (TR: 3000ms) with a different method (permutational entropy) arrived at a similar conclusion (Wang et al., 2017). We have previously detected greater amplitude variation of the MREG signal (Tuovinen et al., 2020) in groups of AD and frontotemporal dementia patients, but not in patients with schizophrenia, thus offering some differential diagnosis capability for fMRI (Tuovinen et al.,2024). We have also found reversed impulse propagation in AD patients as well as increased propagation speed of the cardiovascular impulse mostly in parenchymal areas, which suggests the presence of narrowed and rigid brain arteries (Rajna et al., 2021). Both findings indicated altered impedances of the arterial vessel walls (Mynard et al., 2020), perhaps in relation to reduced perfusion and neurovascular coupling (Solis, et al., 2020) and increased arterial stiffness (Hughes et al., 2014) indicative of reduced elasticity. The current findings of reduced MREG_RHP_ in AD mainly overlap with brain regions having increased pulse speed. The relatively monotonous pulse waveforms in the AD group suggest that their narrowed and inelastic arteries act more like rigid tubes, incapable of reacting to regional physiological blood flow control or precapillary sphincters (Mynard et al., 2020; Rajna et al., 2021).

In line with dysregulated cerebrovascular pathology, the parenchymal resting state networks in mild cognitive impairment and AD patients were reduced in proportion to the severity of the individual’s memory impairment (Rombouts et al., 2005). Indeed, the classical BOLD responses to neurovascular coupling were likewise altered in AD (Buckner et al., 2000; D’Esposito et al., 2003; Rombouts et al., 2005), as an inevitable correlate of cortical atrophy occuring along with changes in neurovascular coupling and blood flow (Johnson et al., 2000). A meta-analysis of positron emission tomography studies with the glucose analogue [^18^F] FDG and fMRI studies showed that AD patients have consistently lesser functional activity than controls in several brain regions, including the medial temporal lobe and frontal pole, during encoding and retrieval cognitive challenge paradigms (Schwindt and Black, 2009).

The typically narrowed and amyloid-laden arterioles in AD patients with cerebral amyloid angiopathy (Weller et al., 2009) fail to dilate and increase cerebral blood flow during systole within normal physiological cardiohemodynamic timeframe. The more monotonous pulse outline in the cardiovascular pulsations reflects reduced accommodation of the stiffened and narrowed cortical arterioles to meet metabolic demand during increased neuronal activity and other factors that would increase regional blood flow. Indeed, the cardiovascular impulses inherently reflect neuronal activation; the brainstem respiratory centers present elevated cardiovascular impulses during the initation and ending of a breath hold task (Raitamaa et al., 2019). Furthermore, the envelope over cardiohemodynamic impulses denoting vascular dilatation precedes by approximately 1.3 seconds the venous BOLD responses in visual cortex upon activation in checkerboard visual stimulation task (Huotari et al., 2022). The loss of vascular responsiveness to sudden changes in regional neuronal activity is deemed to lead to a monotonous fMRI signal arising in association with cognitive impairment.

Another factor that could reduce cardiovascular responsiveness in AD is a failure in the recently described paravascular glymphatic solute convection pathway, which is a contributing factor in the β-amyloid accumulation in periarterial brain tissue in AD (Nedergaard and Goldman, 2020). With declining arterial flexibility and pulsatility, the glymphatic system loses its driving force in the periarterial CSF spaces, which may present a feed-forward mechanism for progressive amyloid accumulation. There is even evidence for reversed impulse propagation in the narrowed and rigid AD vasculature (Rajna et al., 2021), which could induce local damage to blood vessel structures and disrupt the blood-brain barrier (BBB). Indeed, chronic increases in BBB permeability occur at an early stage of AD progression (irrespective of β-amyloid or tau accumulation) in hippocampal areas manifesting and inverted pulse propagation (Nation et al., 2019). We speculate that failed pulsations would further impede convection in a vicious cycle leading to exacerbation of vessel wall and interstitial inflammation, increased protein aggregation, and further failure of arteriolar walls to accommodate physiological cues.

## Summary

In this study we propose a novel, fast and simple FFT-based metric of relative harmonic power (RHP) for quantifying cardiovascular pulse shape in human brain tissue from ultrafast fMRI data capable of critically sampling cardiovascular pulsations without aliasing. We showed that a wide 0.6 – 5 Hz window using flip angle 5° data performs better that with flip angle 25° data. The RHP analysis showed that the cardiovascular pulsations were more monotonous in the AD brain than in controls suggesting a possible failure of neurovascular coupling and responsiveness to physiological demands in brain regions shown in previous research to have increased pulse propagation due to narrowing of (peri)vascular conduits of the glymphatic fluid clearance pathway.

## AUTHOR CONTRIBUTION

Conception and design of the study: AK, VKi

Data Acquisition: NH, VKo, AK, AR, VPo, HH, JT, LR, TV, JKr, VKi, JKa

Analysis: AK, VPe, MA, TV, VKi, JKa

Interpretation: AK,VPe, VKi, JKa

Writing: AK, VPe, MA, VPo, VKo, MJ, NH, HH, JT, JKr, VK, JKa

All authors have contributed to this draft. All authors have approved the final version of this manuscript. All authors have agreed to be accountable for all aspects of this work in ensuring that questions related to the accuracy or integrity of this work are appropriately investigated and resolved.

## ETHICS STATEMENT

This study adheres to the Declaration of Helsinki and institutional approval was granted by the Ethical Committee of Northern Ostrobothnia Hospital District, Oulu University Hospital, currently known as “The regional medical research ethics committee of the Wellbeing services county of North Ostrobothnia”. Written informed consent was obtained from all subjects.

## FUNDING

Research Council of Finland (TERVA funding 314497; TERVA funding 335720; Project 338599 funding, VKi)

Instrumentariumin Tiedesäätiö (*Instrumentarium Science Foundation*) (JKa, TV, JT) Jane ja Aatos Erkko Foundation grant 2016-2021 and 210043 (VKi)

Maire Taponen Foundation (JKa, LR) Medical Research Center Oulu (JKa)

Orionin Tutkimussäätiö (*Orion Research Foundation)* (JKa, VPo)

Suomen Aivosäätiö (*Finnish Brain Foundation*) (JKa, LR, JT, HH) Suomen Lääketieteen Säätiö (*Finnish Medical Foundation*) (JKa, VPo) State research Funding (JKr, JKa)

Emil Aaltosen Säätiö (*Emil Aaltonen Foundation*) (LR, HH) Tauno Tönningin Sääti (*Tauno Tönning Foundation*) (LR)

Juhani Ahon Lääketieteen Tutkimussäätiö sr. (*Juhani Aho Foundation for Medical Research*) (LR) Uulo Arhion muistorahasto (*Uulo Arhio Foundation*) (LR)

The Finnish Parkinson Foundation (HH) Paulo Foundation (HH)

Oulun Lääketieteellinen tukisäätiö (HH)

## CONFLICT OF INTEREST

J. Krüger has served on the advisory board of Novartis, Nutricia, EISAI, Lilly and Roche and received honoraria from lectures from Bioarctic and Lilly and received support for congress participation from Merck.

Other authors have no competing interests to declare

## DATA AVAILABILITY

The data that support the findings of this study are available from the corresponding author upon reasonable request.

## ACKNOWLEDGEMENTS

The authors wish to acknowledge CSC (IT Center for Science, Finland) for its computational resources. The authors also wish to thank Annastiina Kivipää and Matti Pasanen for data acquisition, Jussi Kantola for computational assistance, and Prof. Paul Cumming for critical reading of the manuscript.

## Notes

### Competing Interest Statement

J. Kruger has served on the advisory board of Novartis, Nutricia, EISAI, Lilly and Roche and received honoraria from lectures from Bioarctic and Lilly and received support for congress participation from Merck. Other authors have no competing interests to declare

